# Quantome: A Quantum Surrogate Model for Biophysical Landscapes

**DOI:** 10.1101/2025.08.06.668871

**Authors:** Ashar J. Malik, David B. Ascher

**Author notes:** Correspondence to Ashar J. Malik, David B. Ascher.

## Abstract

Accurately modelling the potential energy landscapes that govern molecular interactions is a central challenge in computational biophysics. While quantum computers promise to solve such problems with high fidelity, a key bottleneck is the encoding of complex spatial information into low-qubit Hamiltonians suitable for near-term devices. Here, we introduce a generalisable framework for creating quantum surrogate models of 2D biophysical landscapes. Our method translates a discrete, classically-derived potential energy grid into a continuous quantum Hamiltonian by fitting it to a high-degree polynomial, where the polynomial’s coefficients directly define the potential energy operator. We demonstrate this pipeline on a custom-designed landscape featuring two asymmetric potential wells. By systematically varying a kinetic hopping term in the Hamiltonian, our variational quantum eigensolver simulations, averaged over 100 runs, successfully reproduce the physical transition from a localised ground state to a delocalised state governed by tunnelling. The entire framework is made accessible through the Quantome web application, a pedagogical platform that allows users to both design custom landscapes and explore pre-calculated results on EF-hand protein binding sites, which serve as a simplified real-world test bed for this framework. The web app is freely available at: https://biosig.lab.uq.edu.au/quantumlabs/quantome.

## Introduction

How do molecules recognise and respond to one another? The answer lies in their physical interactions, a complex interplay of forces and shapes that dictates the machinery of life. These interactions unfold across intricate surfaces, which are best understood as complex potential energy landscapes sculpted by a topology of electrostatic charge and steric hindrance effects [1]. The specific features of this landscape, its energetic valleys of attraction and hills of repulsion, ultimately determine the specificity and strength of a binding event. Such interactions are fundamental across biology, occurring between an enzyme and its substrate, a therapeutic drug and its target receptor, or the monomers of a self-assembling multimeric protein. Therefore, the ability to accurately model these landscapes and predict the outcome of molecular interactions is a central goal of computational biology, holding the key to designing novel therapeutics [2].

Ideally, quantum algorithms in quantum chemistry would allow us to discard classical approximations altogether. The forces we model with proxies like Lennard-Jones potentials and simplified electrostatics are merely macroscopic consequences of complex quantum-mechanical phenomena. A sufficiently powerful quantum computer could simulate these systems from first principles [3], providing a far more realistic view of the interaction landscape. However, the state-of-the-art in quantum hardware is not yet capable of the fault-tolerant computation required for such a full-scale simulation. We are currently in the Noisy Intermediate-Scale Quantum (NISQ) era, where quantum processors are limited by low qubit counts and high error rates, making them unsuited for the deep circuits required by many foundational quantum algorithms [4].

This reality poses a critical question: while we await future fault-tolerant hardware, can the empirically-derived classical landscapes be recast into a format solvable by NISQ devices? Such an approach would allow us to leverage the unique capabilities of quantum processors on tractable problems today. This is precisely the role of hybrid quantum-classical algorithms, with the Variational Quantum Eigensolver (VQE) being a prominent example [5]. VQE acts as a compromise, using a robust classical optimiser to train a shallow, noise-resilient quantum circuit. By doing so, it leverages the strengths of both computational paradigms. Until a true quantum advantage is realised for full simulations, hybrid methods like VQE represent our most promising path forward for applying quantum computation to meaningful scientific problems.

The versatility of VQE has led to its exploration across a range of challenges in the life sciences. In structural biology, researchers have applied VQE to solve simplified protein folding problems by finding the minimum energy conformation of a polypeptide on a lattice [6]. This principle extends to drug design, where VQE is used in pipelines to insert low-qubit kernels into chemically exact workflows for pro-drug activation and covalent inhibition [7]. These applications all build upon VQE’s foundational capability: calculating the ground state energy of a molecular Hamiltonian, first demonstrated on small molecules [8]. This growing body of work illustrates a clear trend toward using simplified, problem-specific Hamiltonians to make near-term quantum devices useful for biology.

Building on this paradigm, we introduce a generalisable framework for creating low-qubit quantum surrogate models of biophysical interaction landscapes. Our method first represents a surface as a discrete potential energy grid, which is then fitted to a high-degree two-dimensional polynomial. The coefficients of this polynomial directly define a sparse Hamiltonian, effectively translating the classical landscape into a format solvable by nearterm quantum processors. We demonstrate this pipeline on two distinct systems: first, on an idealised, user-configurable grid of amino acids, which we have deployed as an interactive web application to serve as a pedagogical tool and second, on a set of complex pre-calculated landscapes derived from Ca^2+^ EF-hand loop protein-ion binding site to validate the method’s applicability. These data are provided for inspection through the web application. We show that this surrogate model maintains high fidelity to the original classical potential and that sampling the VQE-prepared ground state yields a probability distribution sharply concentrated over the landscape’s primary energetic minimum. The aim of this framework is not to demonstrate quantum advantage, but rather to provide a conceptual bridge between the empirically-derived potentials of classical mechanics and the operational capabilities of quantum algorithms, extending the logic of discrete lattice models to continuous energy surfaces.

## Method

### Generation of 2D potential energy landscapes

To demonstrate and validate our framework, we generated two distinct types of 2D potential energy landscapes: a simplified, user-configurable grid of amino acids, and a more complex surface derived from Ca^2+^ EF-hand loop protein-ion binding site. Both systems and their resulting data are made accessible through a unified, interactive web application developed for this work. The application serves a dual role, providing both a pedagogical platform for constructing idealised landscapes and a data viewer for the biophysical landscapes. Ultimately, both methods yield a 9 by 9 grid where each point is described by two key parameters: a distance (r) representing a repulsive boundary and a charge (*q*_*i*_) representing an electrostatic feature.

**Figure 1.**
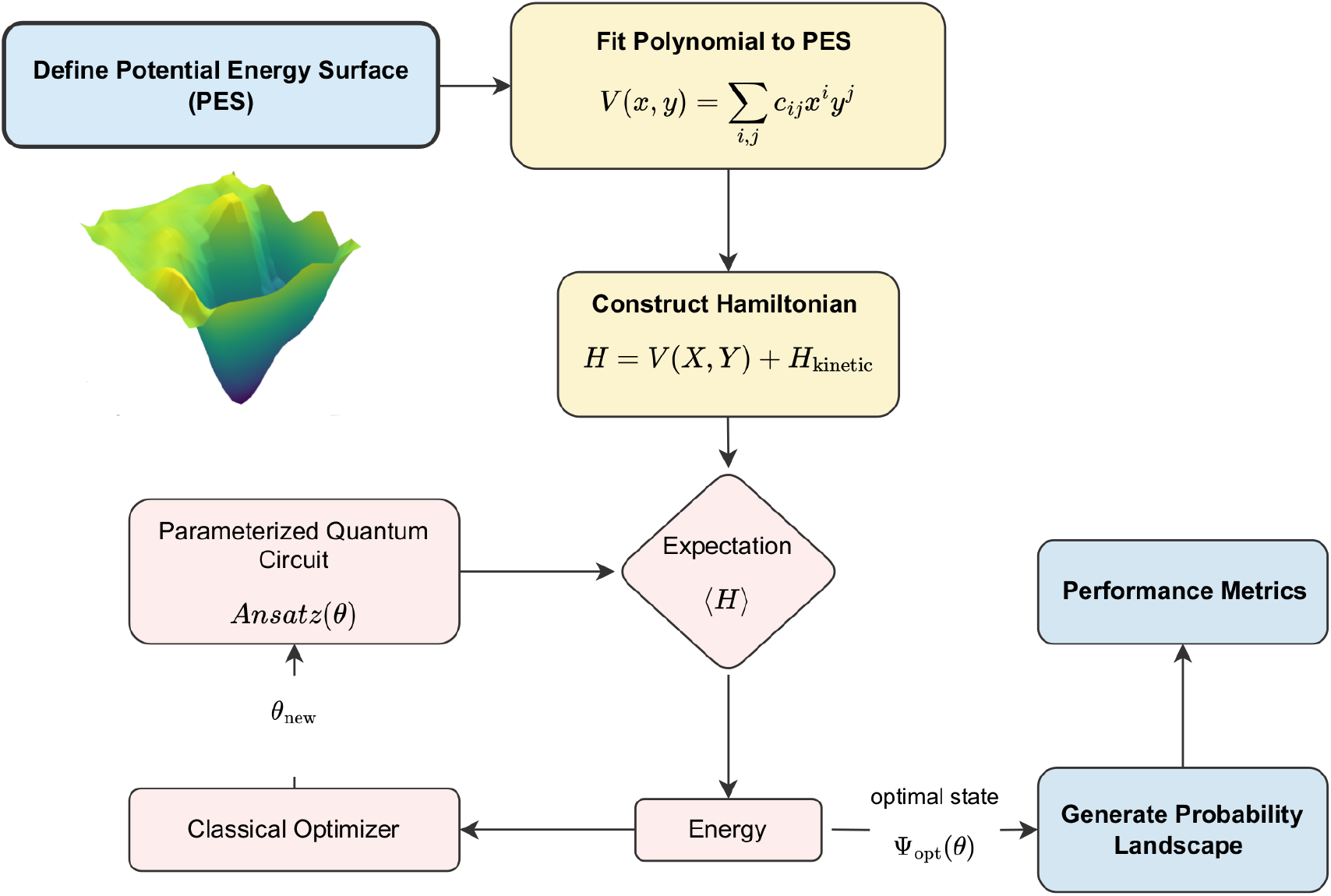
The Quantum Surrogate Model Workflow. The pipeline begins by defining a 2D potential energy surface (PES). This discrete surface is then fitted to a high-degree polynomial to create a continuous surrogate model. The coefficients of this fit are used to construct a matrix Hamiltonian, including a kinetic energy term. The ground state of this Hamiltonian is found using the Variational Quantum Eigensolver (VQE), which uses a classical optimiser to tune the parameters (*θ*) of a quantum circuit (ansatz). The final, optimised state is sampled to generate a probability distribution, which is then analysed using performance metrics.

### User-configurable grid

The idealised grid system allows users to construct a custom landscape by assigning any of the 20 standard amino acids to each point on a 9 by 9 grid. In this idealised model, the lateral distance between adjacent grid sites is not assigned a physical scale and is treated as 1 arbitrary grid unit. For this system, each amino acid is assigned a pre-defined charge and a steric size parameter, which are detailed in the supplementary information. The landscape parameters are determined as follows:

- Distance (*r*): The repulsive distance, *r*, is derived from two user-controlled global parameters: a Probe Plane Height (*h*_*probe*_) and a Minimum Allowed Distance (*ϵ*). Each amino acid is assigned a fixed steric bulk (*s*_*aa*_) representing its size. For any given grid point, the distance is calculated as the separation between the probe plane and the top of that amino acid’s bulk:

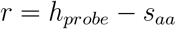

A steric clash constraint is strictly enforced, requiring *r* ≥ *ϵ* for all points on the grid. Therefore, for a calculation to proceed, the user must set a probe plane height, *h*_*probe*_, that is sufficient to clear the bulkiest amino acid present on the grid by at least the distance *ϵ*.

- Charge (*q*_*i*_): The charge parameter at each grid point is the formal integer charge of the user-selected residue.

### Biophysical landscape from protein structure

To generate realistic test cases, we applied our pipeline to protein structures containing EF-hand motifs sourced from the Protein Data Bank (PDB) [9] and identified using Structome- TM [10]. For each selected structure, a specific calcium-binding loop was isolated and processed through an automated workflow to establish a consistent frame of reference and derive the landscape parameters.

- Ion-centric alignment: A crucial pre-processing step was performed to standardise the orientation of each binding site. First, a local point cloud of protein atoms surrounding the bound Ca^2+^ ion was defined. Principal Component Analysis (PCA) was then applied to this point cloud to determine the principal axes of the binding pocket. Based on these axes, the entire protein structure was translated and rotated to place the Ca^2+^ ion at the origin and align the primary exit vector of the pocket with a Cartesian axis.
- Grid definition: Following alignment, a discrete 9 by 9 sampling grid was defined on a plane positioned at a fixed clearance distance from the origin along the aligned axis. The grid was positioned at a clearance of 8.0 Å from the ion and was defined with a 1.0 Å spacing between adjacent points, creating an 8.0 Å x 8.0 Å patch. This grid represents the 2D space from which the protein surface is probed.
- Parameter derivation via ray-casting: From each point on this grid, parameters were derived using a ray-casting algorithm [11].

‐ Distance (*r*): A ray was cast toward the protein. The Euclidean distance from the grid point to the first atom it intersects defines the distance parameter *r*.
‐ Charge (*q*_*i*_): The formal integer charge of the residue to which the intersected atom belongs is assigned as the charge parameter *q*_*i*_.

### Combined potential energy calculation

For both the idealised grid and the biophysical landscape, a potential energy value (*V*) was calculated for each of the 81 grid points. The calculation at each site used the locally-derived distance (*r*) and charge (*q*_*i*_) parameters in a simplified potential function, which combines a Lennard-Jones (LJ) repulsive term with a Coulombic interaction term:

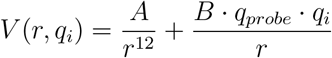

The first term models a hard-sphere steric repulsion, while the second term models the electrostatic interaction between a defined probe charge (*q*_*probe*_) and the local surface point charge (*q*_*i*_). The constants *A* and *B* are global scaling factors for the LJ and Coulombic terms, respectively.

The specific parameters for this function were set according to the system being modelled. For the idealised grid system, the user can set the probe charge *q*_*probe*_ to either +1 or -1, and the scaling factor is *B* = 1. For the biophysical landscapes, which model a Ca^2+^ ion, the probe charge was fixed at *q*_*probe*_ = +2 and a physically-derived scaling factor of 

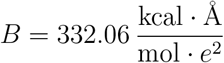

was used. The resulting 9 by 9 matrix of these individual V values constitutes the final discrete potential energy landscape used for all subsequent steps.

### The polynomial surrogate model

To construct a Hamiltonian from the discrete potential energy landscape, it was first necessary to create a continuous analytical function. This was achieved by developing a polynomial surrogate model through a two-step fitting process.

- Surface Interpolation: The discrete 9 by 9 potential energy landscape, V, generated in the previous step was first upsampled to a denser 16 by 16 grid. This was performed using a bivariate spline interpolation (scipy.interpolate.RectBivariateSpline), resulting in a smoother surface that provides a better basis for a robust polynomial fit.
- Polynomial Fitting: A 10th-degree, two-dimensional polynomial, P(x,y), was then fitted to this interpolated 16 by 16 surface. The polynomial takes the general form:

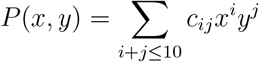

The coefficients, *c*_*ij*_, were determined. This polynomial function, P(x,y), serves as the final, continuous surrogate model for the original discrete landscape. A 10th-degree polynomial was found to provide a sufficient balance between capturing the landscape’s complexity and avoiding excessive oscillations.

- Fidelity Assessment: The accuracy of this surrogate model was quantified by calculating the Root Mean Square Error (RMSE) between the values predicted by the polynomial and the corresponding values on the interpolated 16 by 16 surface.

### Hamiltonian construction

The continuous polynomial surrogate model, *P* (*x, y*), was translated into a matrix Hamiltonian, *Ĥ*, for a corresponding 8-qubit system representing the 16 × 16 grid (2^8^ = 256 states). This was achieved by defining operators for position and energy within this 256-dimensional Hilbert space.

### Position operator representation

The spatial coordinates of the grid were encoded using position operators, *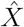* and *Ŷ*. These operators were constructed as 256 × 256 diagonal matrices where the diagonal entries correspond to the x and y coordinates of each of the 256 grid points. Following standard practice for defining operators on a tensor product space, they were formed using the Kronecker product:

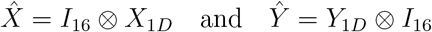

where *X*_1*D*_ and *Y*_1*D*_ are 16 × 16 diagonal matrices of the grid coordinates and *I*_16_ is the 16 × 16 identity matrix.

### Potential and kinetic energy terms

The Hamiltonian was defined with two components: a potential energy term, *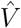*, derived from the surrogate model, and a kinetic energy term, *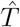*, to allow movement between grid sites.

#### Potential term 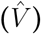

The potential term was constructed directly from the polynomial coefficients (*c*_*ij*_) obtained in the previous step, creating a matrix representation of the potential energy surface:

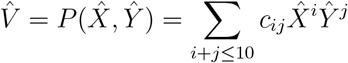

This results in a diagonal matrix where each entry represents the potential energy at the corresponding grid site.

#### Kinetic term 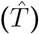

To allow a particle to move between grid sites, a nearest-neighbour hopping term was added. This term introduces off-diagonal elements with a value of −*t*_*hop*_ between adjacent grid points, effectively creating a 2D tight-binding model.

### Final Hamiltonian formulation

The final Hamiltonian is the sum of these two terms, *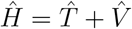*. This complete 256 × 256 matrix was then converted into a SparsePauliOp object, the standard operator format required for execution with Qiskit’s algorithms [12].

### Variational Quantum Eigensolver (VQE) simulation

The ground state of the constructed Hamiltonian, *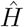*, was determined using the VQE algorithm, implemented with IBM’s Qiskit library [12] making use of a simulator. The VQE workflow consisted of three main parts: algorithm setup, ground state optimisation, and state sampling.

### VQE algorithm setup

The VQE comprised the following:

- Ansatz: A hardware-efficient ansatz, `EfficientSU2`, was chosen as the parameterised trial wavefunction, |*ψ*(*θ*)⟩. The circuit was configured for an 8-qubit system with a repetition depth of 8 `(reps=8)`.
- Optimiser: The circuit parameters, *θ*, were trained using the Simultaneous Perturbation Stochastic Approximation (SPSA) optimiser, a gradient-free method well-suited for the optimisation landscapes of variational algorithms, with a maximum of 1000 iterations.

### Ground state optimisation

The VQE algorithm iteratively finds the optimal parameters, *θ*_*opt*_, that minimise the expectation value of the Hamiltonian:

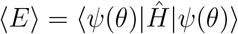

This hybrid quantum-classical optimisation loop was managed using a Qiskit `Estimator` primitive to evaluate the energy at each step.

### State sampling and measurement

Upon convergence, the optimal parameters, *θ*_*opt*_, were assigned to the ansatz circuit to prepare the final ground state approximation, |*ψ*(*θ*_*opt*_)⟩. To determine the probability of finding the particle at each grid location, this final state was measured in the computational basis 16,384 times (`shots=16,384`) using a Qiskit `Sampler` primitive. This process yields a probability distribution over the 256 basis states, which directly correspond to the points on the 16 × 16 grid.

### Analysis and performance metrics

To quantitatively evaluate the VQE simulation results, the final probability distribution was analysed. The specific metrics applied were dependent on whether the system was a biophysical landscape or a user-defined grid.

### Probability distribution mapping

As a first step for both systems, the probability distribution over the 16 × 16 fine grid was downsampled to the original 9 × 9 coarse grid. This was achieved by mapping each point on the fine grid to its nearest-neighbour on the coarse grid and summing the corresponding probabilities.

### Performance evaluation

The resulting 9 × 9 probability distribution was then analysed differently for each system type.

- For the biophysical landscapes, where the centre of the grid represents a known target location (the Ca^2+^ ion site), a full suite of metrics was calculated to assess the performance of the ground state search:
  ‐ **Locational Error (Å):** The Euclidean distance between the center of the grid and the grid point with the highest measured probability.
  ‐ **Enrichment Factor:** The ratio of the probability at the peak location to that of a uniform background.
  ‐ **Probability Mass:** The total summed probability within a 3.0 Å radius of the center of the grid.
- For the user-defined grid system, the analysis focused on visualising the outcome. The final 9 × 9 probability distribution was displayed as a heatmap, and only the **Enrichment Factor** was calculated to quantify the focus of the resulting distribution.

### The Quantome web app

An interactive web application was developed to provide a graphical user interface for the framework presented. The application was built using a Flask (Python) backend to handle calculations and a standard HTML/JavaScript frontend. Molecular visualisation of the protein structures and the overlaid sampling grids is rendered using the Mol* viewer [13], while all 3D interactive plots of the landscapes are generated with the Plotly JavaScript library. The application supports two modes of operation: it loads pre-calculated data for the EF-hand proteins, and for the user-configurable grid, a session management system handles the submission of user-defined landscapes for VQE training. Upon submission, a unique Job ID is issued, allowing the user to retrieve the results once the calculation is complete. The entire application is containerised using Docker and served via an Nginx web server.

## Results

To demonstrate our framework’s ability to probe quantum phenomena, we designed a custom 9 × 9 grid landscape featuring two proximal asymmetric potential energy wells (Figure 2). Using a negative probe (*q* = −1), a deep 2×2 well and a shallower 2×1 well were created with arginine residues (*q* = +1), surrounded by glycine residues. This landscape was processed through our classical pipeline to generate the high-fidelity polynomial surrogate model used for Hamiltonian construction.

**Figure 2.**
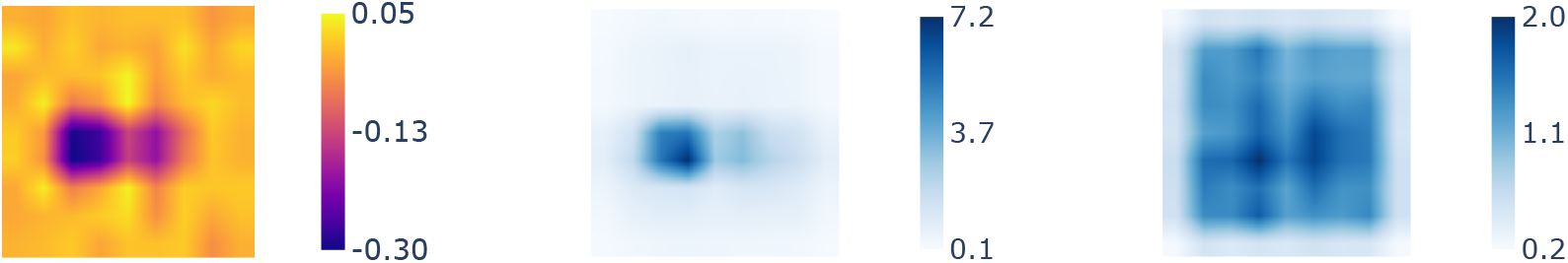
VQE Simulation of a Custom Biophysical Landscape. (Left) A heatmap of the 9× 9 fitted potential energy surface, showing the deep (left) and shallow (right) wells. (Middle) At a low hopping strength (*t*_*hop*_ = 0.01), the averaged VQE enrichment is localised in the deeper well. (Right) At a higher hopping strength (*t*_*hop*_ = 1.0), the ground state delocalises across the grid.

From this single potential, a set of Hamiltonians was constructed, each with a different nearest-neighbour hopping strength (*t*_*hop*_ ∈ {0.0, 0.0001, 0.01, 0.1, 0.5, 1.0, 5.0}) controlling the kinetic energy term. The averaged results from 100 VQE runs for each Hamiltonian reveal a clear transition from localisation to delocalisation. At near-zero hopping strengths (*t*_*hop*_ ≤ 0.1), the enrichment distribution is sharply localised in the deeper of the two potential wells, whose four sites maintain a combined enrichment factor remaining above 20× compared to the background (See supplementary data). As the hopping strength increases, the ground state begins to delocalise significantly. This is seen as a sharp drop in the deep well’s enrichment, as probability spreads across the entire grid. At high hopping strengths (*t*_*hop*_ ≥ 1.0), this delocalisation becomes dominant; indicating that the kinetic energy has overwhelmed the potential landscape.

## Discussion

Potential energy landscapes serve as the fundamental blueprint for molecular interactions, defining the energetically favorable regions that guide binding and recognition. In this work, we constructed such landscapes using a simplified model, combining a short-range repulsive term with long-range electrostatics to represent a surface’s key physical features. The central innovation of our framework is the translation of this classical grid into a continuous, lowqubit quantum Hamiltonian via a high-degree polynomial surrogate. This approach of discretizing a complex biophysical problem onto a grid is analogous to the use of well-established lattice models in protein folding, where the vast conformational space of a polypeptide is mapped onto a simplified, discrete set of states to make the energy calculation tractable. Crucially, our Hamiltonian formalism then allows for the inclusion of a non-diagonal kinetic energy term, controlled by the hopping strength (*t*_*hop*_), which moves the problem beyond a simple classical minimization and allows us to directly probe tunnelling effects.

The results obtained in this work serve as a validation of this framework. By applying the pipeline to a system with a known physical narrative, i.e., the transition from localization to delocalisation, we can confirm its correctness. At near-zero hopping strengths (*t*_*hop*_), the VQE simulation correctly identified the classical ground state, with the enrichment distribution sharply localised in the global potential minimum. As the kinetic energy term was increased, our model captured the onset of tunnelling. This manifested as a global delocalisation where enrichment began to spread across the entire grid, rather than being confined only to the adjacent shallow well. Finally, at high hopping strengths, the framework correctly demonstrated that the kinetic-energy-dominated, where enrichment becomes broadly distributed. This successful reproduction of a well-understood physical transition confirms the consistency and robustness of our surrogate model approach.

While our framework successfully reproduces key quantum phenomena, it is important to acknowledge its limitations. The potential energy model used in this work, particularly in the user-designed grid, is a simplification of a true biophysical landscape. The use of classical Lennard-Jones and Coulombic terms with integer charges serves as a first-order approximation, capturing the dominant steric and electrostatic effects but neglecting more subtle quantum-mechanical interactions like polarisation and charge transfer, which we are reserving for future exploration.

Furthermore, the polynomial surrogate model, while powerful, is itself an approximation of the initial discrete potential. The polynomial fitting process can introduce its own errors, such as smoothing over sharp features, which we observed during the design of our coarse test landscape. Consequently, any experiment conducted using the Quantome web application is sensitive to these layers of approximation. The results should therefore be interpreted as illustrative of quantum behaviour on simplified landscapes, rather than as precise predictions. To test the applicability of our framework beyond idealised scenarios, we applied it to Ca^2+^ binding in EF-hand motifs. This system was specifically chosen because its binding is primarily driven by electrostatic interactions with negatively charged residues, which aligns well with our simplified potential model and avoids the complexities of explicit side-chain coordination. However, this application introduces its own set of approximations. The method of generating the landscape via ray-casting from a flat, orthogonal grid is a significant simplification of a true binding pocket, which is a complex, three-dimensional, and often concave surface.

This approach means our height and charge maps are an approximation of the true steric and electrostatic environment. We posit that these inherent representational inaccuracies are a primary source of the small locational errors observed in our VQE results for the pre-calculated EF-hand systems. Therefore, while the EF-hand serves as a successful proof-of-concept, the results should be viewed as a validation of the quantum surrogate model pipeline on realistic, albeit simplified, biophysical data.

It is important to state that the goal of this framework is not to demonstrate a quantum advantage for this specific problem. Established classical tools can explore more complex surface representations over an entire protein with greater speed and accuracy. The innovation of our work is instead the development of a framework to translate a classical spatial landscape into a quantum Hamiltonian. This surrogate model serves as a conceptual bridge to probe quantum phenomena on these surfaces. The true promise of such an approach lies in its scalability to problems that are classically intractable. One can conjecture an entire protein’s surface encoded into a single, vast quantum state; a quantum algorithm could then search this exponentially large space to answer questions such as “where would a given ion localise?” Our work represents a foundational step for this vision, developing and validating the principles of spatial encoding and Hamiltonian construction required for such future applications.

The framework presented here is foundational, with several clear avenues for future development. The physical realism of the potential energy landscapes can be improved by incorporating more sophisticated classical force fields that include effects such as partial charges and polarisability. Methodologically, extending the landscape generation and Hamiltonian construction to three dimensions is a key next step. Beyond finding static ground states with VQE, a compelling direction is to use these surrogate Hamiltonians to simulate quantum dynamics, allowing for the study of time-dependent phenomena like the evolution of a wavepacket during a binding event. The underlying framework provides a modular base for developers to prototype these future extensions, while the deployed Quantome web application serves as an accessible pedagogical platform for the wider community to build intuition for how quantum algorithms interact with classical energy landscapes.

## Conclusion

In this work, we have introduced and validated a novel framework for creating low-qubit quantum surrogate models of biophysical landscapes. By translating a discrete, classical potential energy surface into a continuous polynomial, we constructed a Hamiltonian that successfully modelled the transition from a localised state to a delocalised one governed by tunnelling. Our simulations demonstrate the viability of this approach for probing quantum phenomena on complex surfaces. This work, together with the accompanying, freely available, Quantome web application, provides both a foundational method and an accessible tool for exploring the interface of classical biophysics and near-term quantum computation.

## Code and Data Availability

The Quantome web application is freely available at: https://biosig.lab.uq.edu.au/quantumlabs/quantome

