## Supplementary Information for "Quantome: A Quantum Surrogate Model for Biophysical Landscapes"

### Supplementary Table

Table S1: Simplified steric bulk parameters for the 20 standard amino acids, sorted alphabetically. The formal integer charges used in the potential function are: Asp (-1), Glu (-1), Lys (+1), and Arg (+1). All other unlisted amino acids are treated as neutral.

| **AA** | **Bulk** | **AA** | **Bulk** | **AA** | **Bulk** | **AA** | **Bulk** | **AA** | **Bulk** |
|---|---|---|---|---|---|---|---|---|---|
| Ala | 2 | Arg | 9 | Asn | 4 | Asp | 5 | Cys | 3 |
| Gln | 6 | Glu | 7 | Gly | 1 | His | 7 | Ile | 5 |
| Leu | 5 | Lys | 7 | Met | 6 | Phe | 8 | Pro | 3 |
| Ser | 3 | Thr | 4 | Trp | 10 | Tyr | 9 | Val | 4 |

### Supplementary Figures

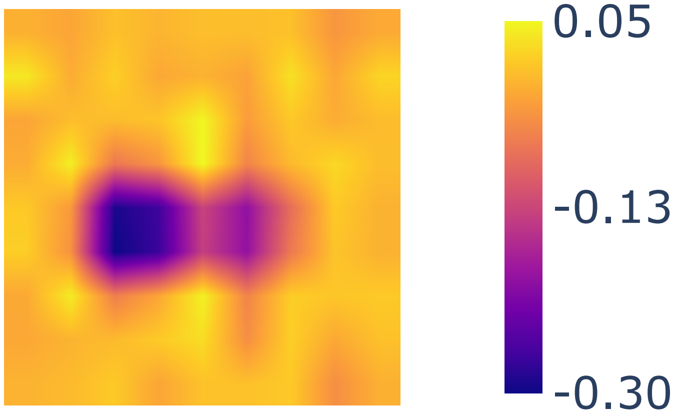


Figure S1: **Fitted Potential Energy Landscape.** Top-down view of the 9x9 potential energy surface, downsampled from the 16x16 polynomial fit. This surface forms the potential term of the Hamiltonians.

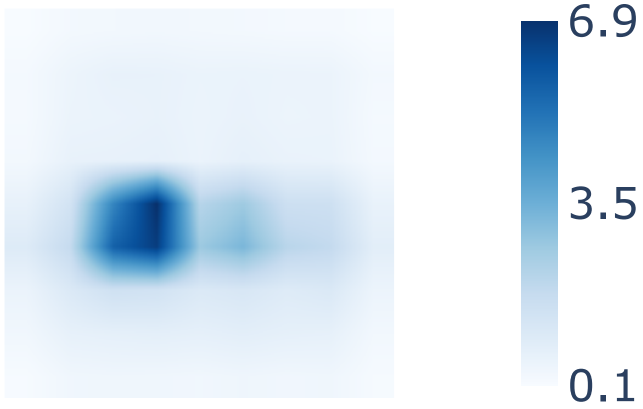


Figure S2: **VQE Enrichment Distribution for $t_{hop} = 0.0$.**

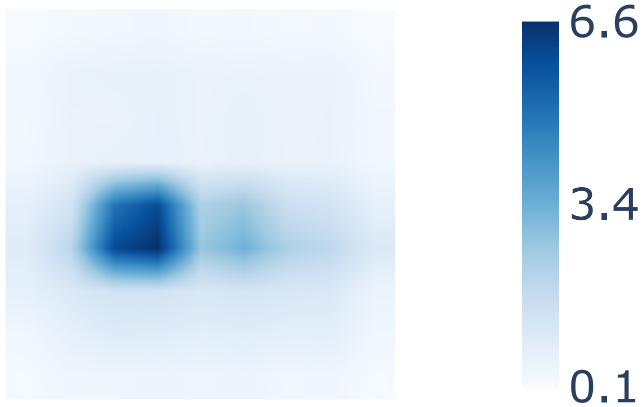


Figure S3: **VQE Enrichment Distribution for $t_{hop} = 0.0001$.**

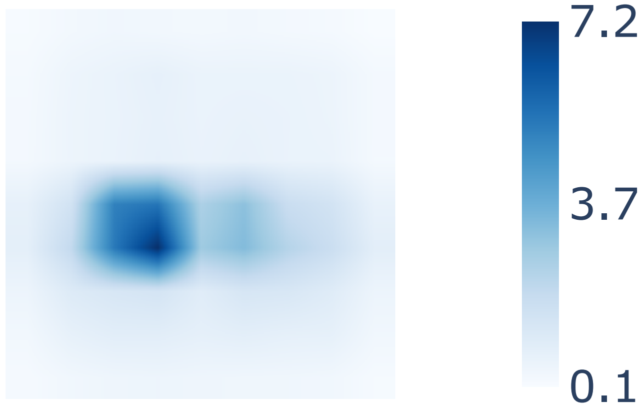


Figure S4: **VQE Enrichment Distribution for $t_{hop} = 0.01$.**

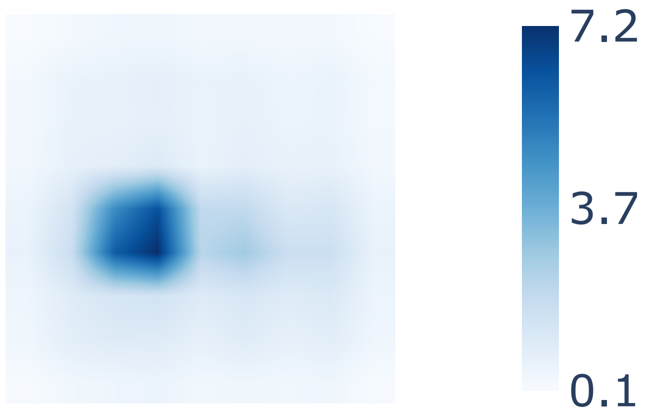


Figure S5: **VQE Enrichment Distribution for $t_{hop} = 0.1$.**

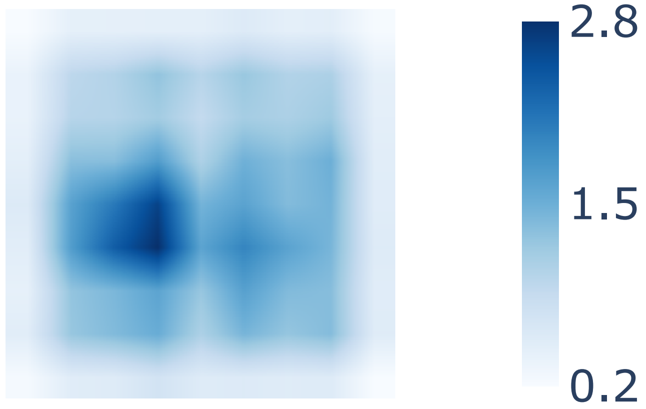


Figure S6: **VQE Enrichment Distribution for $t_{hop} = 0.5$.**

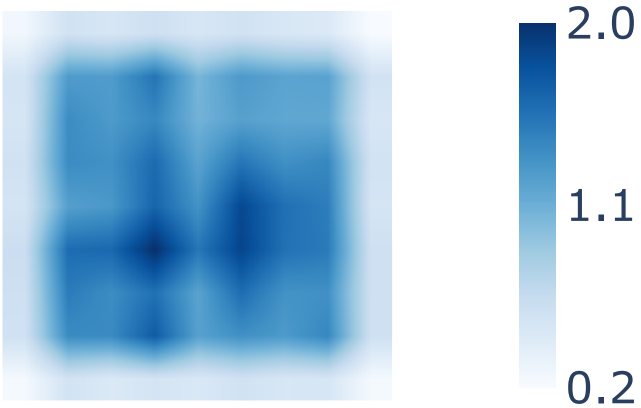


Figure S7: **VQE Enrichment Distribution for $t_{hop} = 1.0$.**

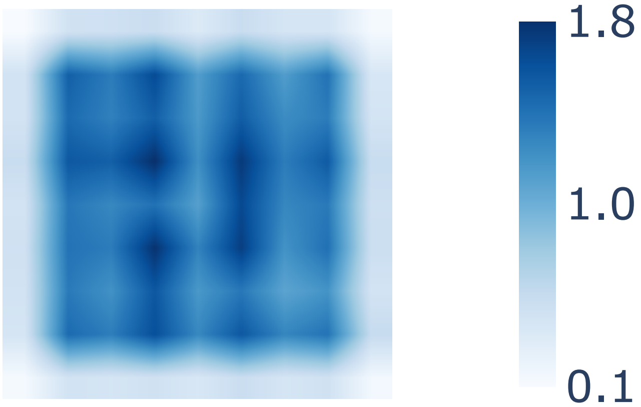


Figure S8: **VQE Enrichment Distribution for $t_{hop} = 5.0$.**

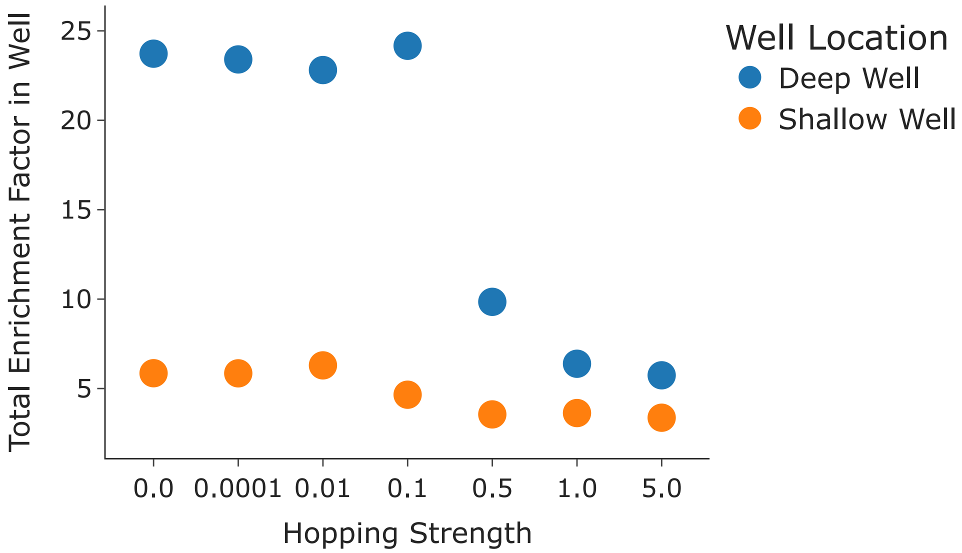


Figure S9: **Quantitative Analysis of Delocalization.** Total enrichment factor within the deep and shallow wells as a function of the hopping strength ($t_{hop}$). The enrichment in both wells decreases at high $t_{hop}$ as the ground state delocalizes over the entire grid.
